# Cortically-evoked movement in humans reflects history of prior executions, not plan for upcoming movement

**DOI:** 10.1101/2022.10.27.514074

**Authors:** Abdelbaset Suleiman, Deborah Solomonow-Avnon, Firas Mawase

**Affiliations:** Faculty of Biomedical Engineering, Technion – Israel Institute of Technology, Israel

**Keywords:** evoked movement, TMS, primary motor cortex, history of movement, default plan, ongoing motor planning

## Abstract

Human motor behavior involves planning and execution, but we often perform some actions more frequently. Experimentally manipulating the probability distribution of a movement through intensive repetition toward a certain direction causes physiological bias toward that direction, which can be cortically-evoked by transcranial magnetic stimulation (TMS). However, because movement execution and plan histories were indistinguishable to date, to what extent TMS-evoked biases are due to more frequently executed movement, or recent planning of movement, is unclear. Here, we use novel experimentation to separately manipulate recent history of movement plans and execution, and probe the effects of this on physiological biases using TMS, and on default plan for goal-directed actions using a behavioral timed-response task. At baseline, physiological biases shared similar low-level kinematic properties (direction) to default plan for upcoming movement. However, when recent movement execution history was manipulated via thumb movement repetitions toward a specific direction, we found a significant effect on physiological biases, but not plan-based goal-directed movement. To further determine if physiological biases reflect ongoing motor planning, we biased movement plan history by increasing the likelihood of a specific target location, and found a significant effect on the default plan for goal-directed movements. However, TMS-evoked movement during the preparation period did not become biased toward the most frequent plan. This suggests that physiological biases provide a readout of the default state of M1 population activity in the movement-related space, but not ongoing neural activation in the planning-related space, potentially ruling out relevance of cortically-evoked physiological biases to voluntary movements.

**Highlights:**

- Stimulating the human motor cortex selectively evoked thumb movements toward a specific direction (physiological bias)
- At baseline, these physiological biases shared similar low-level kinematics with default plan for voluntary goal-directed movements
- Modulating the probability distribution of prior movements had a significant effect on physiological biases
- However, biasing history of plans for upcoming movement toward a specific direction had no effect on evoked movement direction
- During ongoing planning of voluntary movement, evoked movements maintained the distinct and robust baseline bias, regardless of change in probability distribution of history of upcoming plans

## Introduction

Human motor performance depends not only upon the ability of the sensorimotor system to plan and execute movements that are relevant to the current state of the environment and/or the body, but also to the recent history of these actions. One emerging approach to understanding the effects of this recent history on the human brain is through the study of directional biases in involuntary movements evoked by transcranial magnetic stimulation (TMS) applied over the motor cortex^1,2^. TMS provides a unique and powerful measure compared to other techniques, for example, because of the causal activation of effector-specific motor representation that can be precisely evoked and is not easily observed otherwise. For instance, TMS over the primary motor cortex (M1) generally evokes thumb movements in a consistent direction, but when participants voluntarily move their thumbs repeatedly over minutes in an opposite direction, subsequent TMS pulses now evoke thumb movements in the recently practiced direction^1–3^. These physiological, practice-dependent, TMS-evoked directional biases are robust, reliable and last for tens of minutes before returning to the baseline direction^1–3^.

What is the mechanism, at the behavioral and neural level, that drives these profound history-dependent physiological biases in the motor area? One possibility is that these are execution-dependent biases that provide a measurable readout of the physiological changes induced by motor practice^2^ or learning^3^ paradigms. Thus, they might reflect a change in potentiation of synapses that are repeatedly activated in the very recent past^1,4,5^, or changes in the tuning function of neuron population activity in the direction of a repeated action, such that when a stimulation pulse is applied over the neural population within M1, a new “default” state is elicited^6–8^.

Activity in M1, however, is not concerned only with generating motor commands, but is also involved in the processing of higher-order signals for motor planning such as action selection^9–13^. Using non-invasive TMS over M1 as a readout of the functional state of the motor cortex in humans during the planning period (i.e., time prior to an overt action that is unconfounded by descending motor commands), Klein-Flügge and Bestmann (2012)^14^ elegantly showed that the motor system is dynamically shaped by our prior expectations about forthcoming movements. Such prior expectations can be formed by variables that are relevant for action selection, such as the expected reward that can be obtained following an action, or the uncertainty about the required action, given the instruction cue^15,16^. Thus, an alternative explanation is that the evoked physiological biases might also reflect biases toward prior plans of upcoming movements. When hundreds of single-direction movements are repeatedly planned and executed in the recent past, it is possible that neural activity associated with the practiced movements becomes biased, not only toward the most frequently executed movement, but also toward the most probable movement to come next^17^. Although not directly tested in TMS paradigms, this hypothesis was recently supported in goal-directed behavioral experiments by showing that behavioral biases primarily originate from changes associated with prior planning of movement, and to a much lesser extent from changes associated with prior executed movement^17,18^. That is, participants seem to generate a default plan associated with a practiced or recent movement, that can then be modified when the context requires a different action^17^. Similar plan-based directional biases have been reported following repeated passive movement ^3,19^.

Because prior history of movement executions and plans are indistinguishable and largely overlapped in previous work, it remains unclear to what extent the physiological biases evoked by stimulation over the motor cortex are due to effect of pure execution-dependent prior history, and/or to prior planning-dependent history of the most probable action and follows the same plan-based mechanism observed in previous behavioral work. Here we systematically dissociate these possibilities using novel experimental manipulation of recent execution history and plan history, and probe movement execution and plan biases, resulting from the manipulation, using a TMS paradigm and a behavioral timed-response task, respectively. We show that physiological biases reflect a process that is profoundly affected by the recent history of executed movement, with no evidence for a process sensitive to ongoing planning of future actions. We suggest that repetition-induced physiological changes provide a readout of the default state within the movement-related neural space and have little effect on ongoing neural activation related to planning of subsequent voluntary movements. Our findings also raise the possibility that evoked physiological biases and the well-documented behavioral biases reflect dissociable effects likely originating from different neural mechanisms. Lastly, we discuss the potential influence of higher-level perceptual decision-making (e.g., processing sensory information or strategic decision) and motor planning-related processing (e.g., action-selection) on the physiological and plan-based biases we observed.

## Results

### Experiment 1: Baseline physiological biases shared similar low-level movement kinematic properties with plan-based biases

We started with the question of whether the physiological biases evoked by TMS and the plan-based biases at baseline are similar or dissimilar. We predicted that if physiological biases reflect a distinct process from plan-based biases, then the distribution of TMS-evoked movements should show little overlap with the distribution of participants’ baseline default plans. If movement distributions, however, show large overlap at baseline, then at least two explanations might be possible. First, physiological biases may be sensitive not only to history of the executed movement, but also to planning, supporting the hypothesis that these processes might reflect similar underlying mechanism/s. Alternatively, both processes may be distinct, yet converge to a similar state of the neural activity, presumably reflecting a default movement affected by prior hand use, that can be evoked by the TMS stimulation.

We analyzed the direction of the thumb, our experimental end-effector, during a protocol that included a goal-directed behavioral session preceded and followed by a TMS session. In general, in the TMS neurophysiological sessions, single pulses were delivered over the thumb area of the motor cortex that evoked involuntary and isolated thumb movements (Fig. 1A). In the goal-directed behavioral session, participants were asked to make fast voluntary center-out ballistic movements with the same thumb toward different targets presented at an equal distance from the starting point. Thumb-to-cursor mapping was done using the following angle-to-thumb-direction definitions: 0° -adduction, 90° -extension, 180° -abduction and 270 °-flexion. For example, to reach a target at 135°, participants needed to move the thumb in the extension-abduction direction (Fig. 1B).

**Figure 1.**
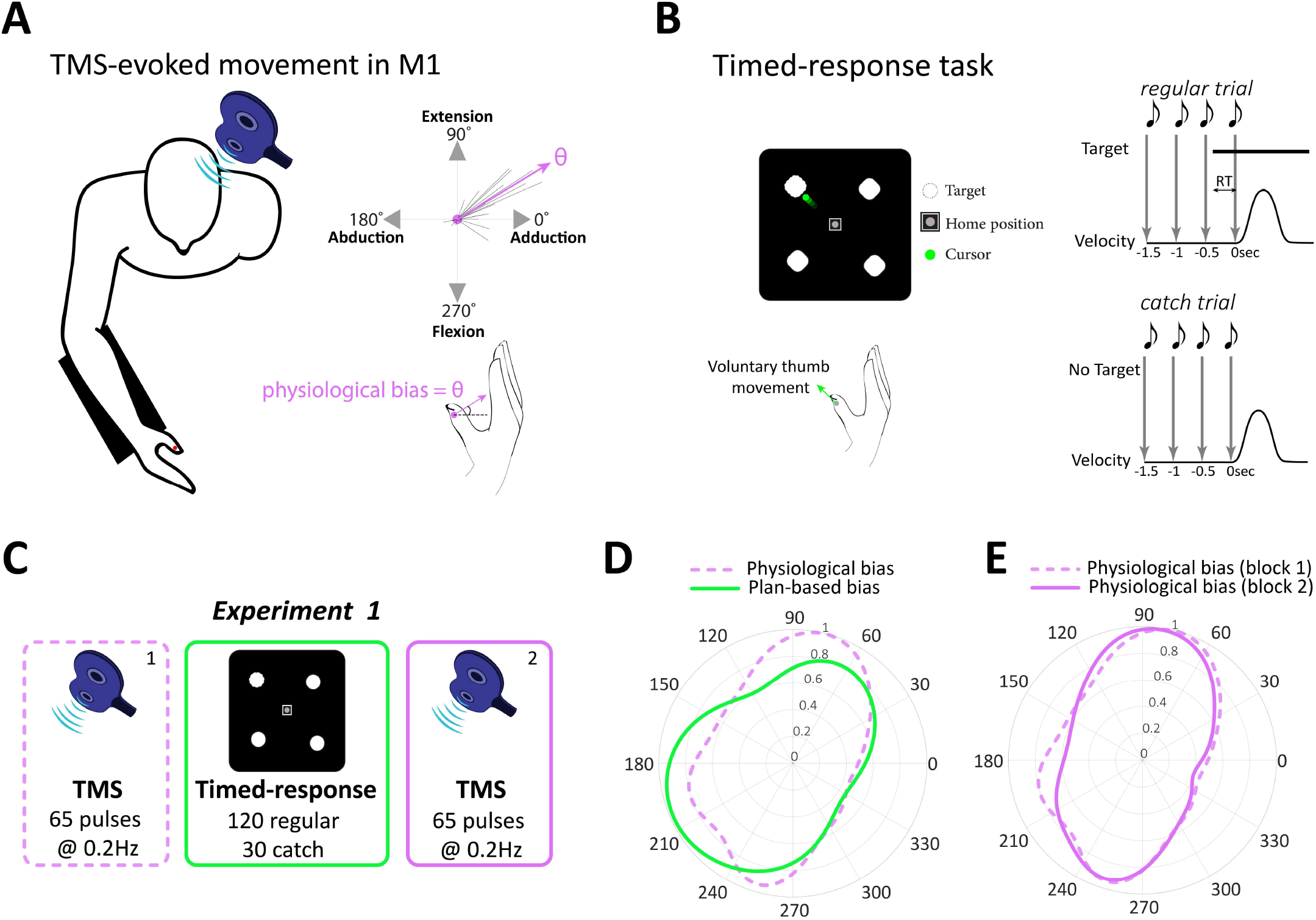
Experimental setup, protocol and results of Experiment 1. **A**. Schematic overview of the TMS protocol. We delivered TMS over the motor cortex to elicit thumb movements and measured physiological bias as the evoked movement with maximum probability. Dashed lines in top right graph represent multiple evoked thumb movements. **B**. Timed-response task. Participants controlled the cursor presented on a monitor by voluntarily moving their thumb. In *regular trials*, participants initiated their movement synchronously with the onset of the fourth tone toward a target (top panel). In *catch trials*, no target ever appeared but participants were still required to plan and move to any direction with the onset of the fourth tone (bottom panel). RT indicates effective reaction time. **C**. Protocol of Experiment 1. Participants performed a block of TMS (65 pulses), followed by a block of goal-directed timed-response tasks (150 trials), and lastly a block of TMS (65 pulses). The 2^nd^ TMS block was conducted to make sure that the relatively short block of voluntary movements (150 trials) did not lead to significant physiological biases. **D**. von Mises probability distribution of physiological biases (i.e., TMS-evoked movements; magenta) and plan-based biases (i.e., green). **E**. Distributions of physiological biases in TMS blocks 1 and 2, indicating that short practice of voluntary movement was not sufficient to induce significant physiological changes. Data in D-E is from all participants pooled together.

In Experiment 1, participants underwent a single TMS block in which 65 pulses were delivered to estimate the physiological biases at baseline, and two goal-directed blocks (before and immediately after the TMS block) in which participants were asked to perform timed-response thumb movements toward a target, selected from four possible targets, each presented randomly 30 times (120 thumb movement in total). In each timed-response trial, participants heard a sequence of four tones, spaced 500 ms apart. The selected target appeared 50-300 ms prior to the fourth tone, and participants were instructed to move to the target as quickly as possible in synchrony with the onset of the fourth tone^20–22^. The predictable sequence of auditory tones serves to minimize the ambiguity about the time of target presentation, which is known to increase reaction times (RTs)^20,23^. We also included a subset of catch trials (30 trials) in which no target ever appeared, and participants were required to move with the onset of the fourth tone to any direction (Fig. 1C, see Methods and Materials). These catch trials enabled us to assess participants’ default plan for upcoming movement in the absence of a target presented in a specific location (see Discussion for possible cognitive-related interpretations of what was measured in catch trials). In total participants performed 150 voluntary thumb movement in the timed-response block.

Distribution of thumb direction (during both evoked and voluntary conditions) was estimated using von Mises distribution^24^. Two measures of biases were then defined: *physiological biases*, defined as the direction with highest probability of the fitted von Mises distribution on the TMS-evoked movements and *plan-based biases*, defined as the direction with highest probability of the fitted von Mises distribution on the plan-based movements during the catch trials of the timed-response block. Our data indicates that physiological biases and plan-based biases at baseline were highly similar, as both distributions showed large overlap (Kuiper’s test, *p* = 0.41, Fig. 1D), suggesting that representation of thumb movement on the motor cortex, as revealed by TMS, shares similar low-level kinematic properties (e.g., direction) with participants’ default plan for impending movement execution during the timed-response trials. The distribution of the default plan directions in catch trials, as well as that of TMS-evoked movement, was not uniform, but was biased toward particular directions, dominated by the far right (30°–60°) and near left (190°–270°; Fig. 1D). These movement directions are consistent with previous work^20,25^ and were likely preferred because they are the least effortful^26^, and/or influenced by thumb use in daily motor functions.

Consistent with previous work^1,27^, we confirmed that short practice (∼8.5 min) of voluntary goal-directed movements during the timed-response block, together with the fact that participants made movements equally to different directions, was insufficient to induce physiological biases, and thus would not confound subsequent measurements. This was revealed by the similar distribution (no significant difference, *p* = 0.91) of physiological biases before (TMS block #1) and after (TMS block #2) the goal-directed block (Fig. 1F). In addition, movements that were inadvertently initiated before, or immediately after the target cue (i.e., too early for the participant to have actually processed the target and planned accordingly; ≤50 msec after target presentation^55^), in the timed-response trials (hereafter termed *inadvertent movements*, meaning movements that were not resulting from or achieved through deliberate planning) were analyzed and compared to catch trials to test whether or not they were essentially equivalent. To test this, we analyzed the direction of all inadvertent movements (10.94% of all trials) occurring any time before the target appeared, and up to 50 msec immediately after the target was displayed. We found that movement direction in these trials was comparable to that in catch trials (no significant differences between distributions, *p* = 0.69) (Supplementary figure S1A). Supporting evidence that the inadvertent-movement directions indeed reflected default plan (i.e., default preparation of upcoming movement in the absence of a presented target) is the fact that the plan for movements to targets that appeared near the default directions should have been readily available and when the target cue comes in, the participant should accurately reach the target with a shorter RT. We thus expected increased accuracy and shorter RTs for a target near the plan-based biases (defined as the closet target with minimal distance (≤ 45°) from the plan-based bias), but not for the other targets. Indeed, fitting the speed-accuracy tradeoff function (SAF, Supplementary Fig. S1B-C) revealed significantly (*p* = 0.042) increased accuracy (as reflected by the parameter *α*_0_ that defined the participant’s lower limit accuracy) at short latencies on targets near the plan-based direction (mean *α*_0_ of 0.368 with 95% CI of [0.037 0.581]), but not near the other targets (mean *α*_0_ of 0.0728 with 95% CI of [0.01 0.169]) (Supplementary Fig. S1D-E).

### Experiment 2: History of prior movements modulated physiological biases, not plan-based biases

Experiment 1 indicated similarity between physiological biases and plan-based biases at baseline. Next, we asked whether manipulating the prior history of executed movement would similarly affect movements evoked by TMS (i.e., physiological biases) and plan-based biases. In Experiment 2, we systematically manipulated the recent history of executed movements by asking participants to repeatedly make thumb movements toward a new direction located in the opposite direction of each participant’s baseline physiological bias. We estimated the physiological bias using TMS, and the plan-based movement bias via the catch trials of the timed-response block, before and after participants (n=15) voluntarily repeated, for 40 min, a movement toward the center (*φ*_*rep*_) of a semi-circular arc target set at 180° from *θ*_*base*_ (Fig. 2A) (see Experimental Procedures).

**Figure 2.**
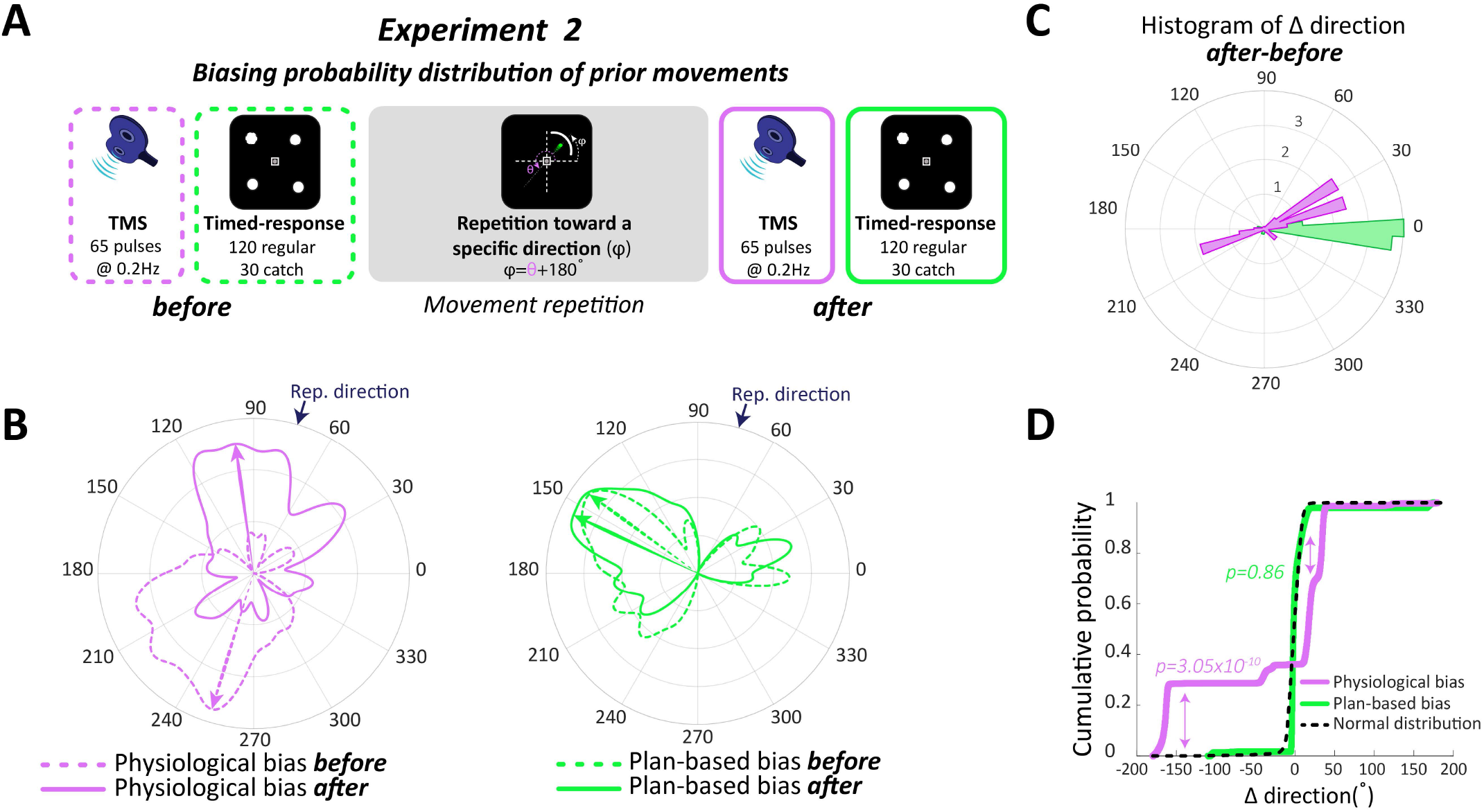
Movement history modulated physiological biases, not plan-based bias of voluntary movements. **A**. Protocol of Experiment 2. **B**. von Mises probability distribution of physiological biases (left) and plan-based biases (right) before (dashed) and after (solid) changing probability distribution of prior movements through repetition toward a novel direction. Data from an individual participant. **C**. Histogram of change (*after*-*before*) in direction of physiological biases (magenta) and plan-based biases (green). **D**. Cumulative distribution function of differences in directions for physiological biases, plan-based biases, and null hypothesis of normal distribution (black dashed line).

We found a significant effect (Kuiper’s test, *p* = 3.18 × 10^−9^) of prior history on physiological biases as seen by the prominent changes in the evoked movement toward the most recently practiced direction (Fig. 2B-D). In contrast, when we probed plan-based bias during the catch trials in the timed-response block, we found no evidence for changes in default plan distribution (*p* = 0.78, Fig. 2B-D). This data suggests that physiological biases are largely affected by execution history. This result may be taken as evidence that repeated movements lead to a change in cortical network representing preferred thumb movements, potentially reflecting a new state in the neural space within the motor cortex to which the neural activity converged into following the changes of recent movement history. In contrast, physiological biases were not dependent on plans of upcoming movements.

Control analysis confirmed that movements that were initiated before, or immediately after, the target cue (i.e., inadvertent movements) were essentially equivalent to default directions in catch trials and were processed accordingly, as in Experiment 1. The distribution of movement directions of all inadvertent movements in the timed-response trials occurring any time before the target appeared, and up to 50 msec immediately after the target was displayed (12.22% of all trials), was comparable to that of the default movements in the catch trials (no significant differences between distributions, *p* = 0.86) (Supplementary Fig. S2A). As expected, and as in Experiment 1, fitting the speed-accuracy tradeoff function (SAF) revealed significant (*p* = 0.011) increased accuracy (as reflected by the parameter *α*_0_ that defined the participant’s lower limit accuracy) at short latencies on targets near the plan-based direction (mean *α*_0_ of 0.531 with 95% CI of [0.429 0.694]), but not near the other targets (mean *α*_0_ of 0.068 with 95% CI of [0.01 0.3718]) (Supplementary Fig. S2B-C).

### Experiment 3: Movement-related physiological changes were distinct from ongoing planning-related processing

The results of Experiment 2 emphasized that while physiological biases were impacted by recent practice, the training had little effect on subsequent movement plans. During catch trials (no target appeared) or regular trials with low RTs (i.e., trials in which the target appeared but participants respond very early before processing the true location of the target(, participants prepared a default plan similar to that at baseline, despite the large post-training biases in movement evoked by TMS. Yet, since repetitive movements in Experiment 2 involved both planning and execution to repeatedly reach the same target, it is impossible to determine whether the observed physiological biases were affected by the repetition of planning, execution, or both. In addition, from the results of Experiment 2, we cannot determine whether TMS evoked movements reflect, to any extent, ongoing planning. In fact, probing of the physiological biases by TMS was done after the timed-response task, and thus outside the context of the timed-response task, when the participants were not planning to make any voluntary movements. In Experiment 3, we addressed these points by systematically dissociating the effect of prior planning of upcoming movements from prior history of executed movements, using a modified Go/No-Go paradigm, with interleaved TMS pulses to probe physiological biases during the movement preparation period. We controlled for the repetition frequency as in Exp. 2, but instead of executing hundreds of movements toward a specific direction, here, in most trials (80% of trials) participants repeatedly prepared a specific plan without executing the movements.

If physiological biases reflect a readout of ongoing activity that is related to planning of subsequent goal-directed movement, in Experiment 3 we would expect a substantial and dynamically increasing bias in the direction of the evoked movement during the preparation period, toward the upcoming movement that participants are preparing for. Alternatively, we predicted that if physiological biases do not reflect a readout of ongoing planning-related processing ahead of upcoming movement and have little relevance with respect to goal-directed voluntary behavior, changing the distribution of potential targets should affect only the plan-based biases, but not the TMS-evoked physiological biases.

To test this, we biased the probability of upcoming movement plans by making potential target locations in the timed-response task not equally probable and, instead, increased the likelihood of targets presented at specific locations in the workspace (Fig. 3A). Unlike the baseline block in which presentation of the four targets was uniformly distributed, here, the most frequent target was presented in 70% of trials (525 trials), and the other three targets were each presented in 10% of trials (75 trials each target), with a total 750 trials. To lead participants to repeatedly plan a movement without actually executing it, we used a modified delayed reaching Go/No-Go paradigm. Here, in 80% of trials participants were led to prepare a movement but did not execute it (No-Go condition), and in 20% of trials, participants executed their planned movement (Go condition) (see Methods and Materials). Importantly, to test the relationship between physiological biases and ongoing planning with respect to goal-directed voluntary behavior, single TMS pulses were delivered in random trials after the target was presented but 150 msec before the Go/No-Go cue, allowing for sufficient time to prepare the cued movement prior to delivery of TMS (Fig. 3B).

**Figure 3.**
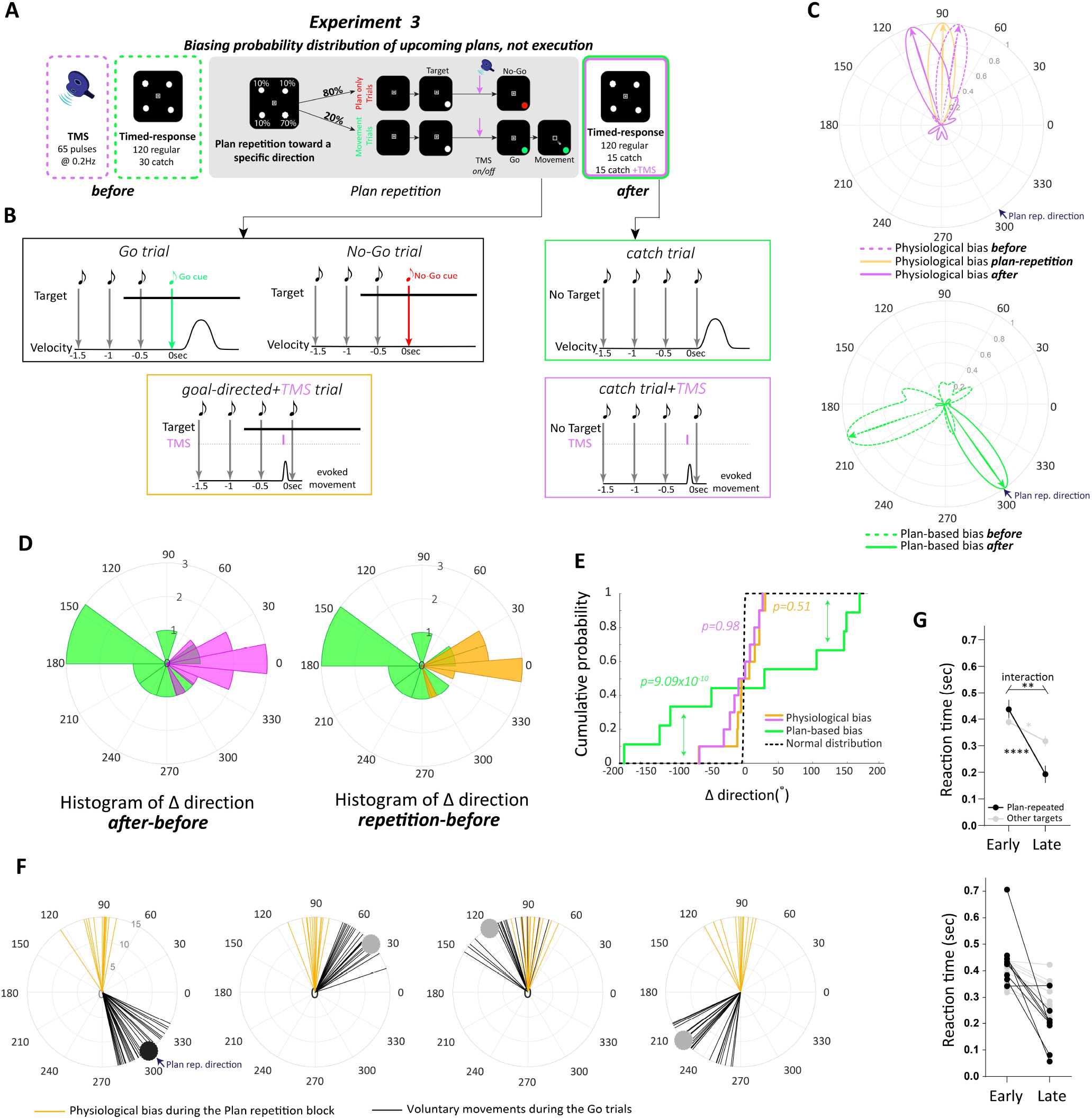
Movement-related physiological changes were distinct from ongoing planning-related processing. **A**. Protocol of Experiment 3. **B**. Trial schedule of each trial type. During Go/No-Go trials, the target appeared 750msec before the cue (target turned green in Go trials while in No-Go trials the target turned red). In some trials, after presenting the target, a single TMS pulse was delivered 150 msec before the Go/No-Go cue. This allowed sufficient time to prepare the desired movement. During the post-plan-repetition block, we implemented the timed-response task that including some catch trials where target never appeared. In addition, in some of these catch trials, a single TMS pulse was delivered 150 msec before the 4^th^ tone. **C**. von Mises probability distribution of physiological biases (top) before (dashed magenta), during (solid orange) and after (solid magenta) the plan repetition block. von Mises probability distribution of plan-based biases (bottom) before (dashed green) and after (solid green) the plan repetition block. Data from an individual participant. **D**. Histogram of change in direction of physiological biases (*after*-*during plan repetition:* orange, *after*-*before:* magenta), and plan-based biases (green). **E**. Cumulative distribution function of differences in directions for physiological biases, plan-based biases, and null hypothesis of normal distribution. **F**. Thumb movements (black lines) during the Go trials and TMS-evoked thumb movement during ongoing planning of the desired movement (orange lines). Each circular panel represents the distribution of movement for each of the four targets during the plan repetition block. Data suggests robust and invariant physiological biases regardless of the plan. **G**. Reaction time (sec) for movements in the Go trials toward the most frequent target (black) and the other targets (gray). Early and late represent the first and last 10 trials of the repetition block, respectively. Average across participants (top) and individual data are shown (bottom). Data show mean ± s.e. with *p<0.05, ** p<0.01 and **** p<0.0001.

Our data revealed that repetition of a plan for a specific movement had no significant effect on physiological biases. When participants repeatably planned a movement toward a specific direction, TMS-evoked movement direction did not change, but instead revealed robust biases toward the baseline direction regardless of ongoing plan (Fig. 3C, top panel). The cumulative distribution function of difference, comparing the distribution of TMS-evoked movement during the plan repetition block with the baseline block, showed non-significant difference (Kuiper’s test, *p* = 0.51) with the null hypothesis of normal distribution (*µ* = 0, *σ* = 10°) (Fig. 3D-E). However, in contrast, changing the likelihood of potential targets had a significant effect on planning of upcoming movements (Fig. 3C, bottom panel). The cumulative distribution function of difference in plan-based directions, during the catch trials, showed significant shift (Kuiper’s test, *p* = 9.09 × 10^−9^) from the normal distribution, toward the most frequently planned target (Fig. 3D-E).

To further confirm that TMS-evoked physiological biases were indeed distinct from any activity related to ongoing planning, we applied single TMS pulses in some catch trials at 150 msec before the 4^th^ tone in the final timed-response block after plan repetition (Fig. 3A-B). This test allowed direct investigation of the relationship between physiological biases and the newly altered default plan for voluntary movement. Our data revealed consistent results; despite profound biases in the default plan toward the most frequent plan, TMS-evoked movement showed robust biases toward the baseline direction, indifferent to the new plan-based biases (Fig. 3C). The cumulative distribution function of difference, comparing the distribution of TMS-evoked movement after the plan repetition block with the baseline block, showed non-significant difference (Kuiper’s test, *p* = 0.98) with the null hypothesis of normal distribution (*µ* = 0, *σ* = 10°) (Fig. 3D-E). Figure 3F shows individual data of evoked movements (orange lines) during the plan repetition block when the participant was preparing and just before making a goal directed movement toward the different targets. These elicited movements were largely different from the actual movements (black lines) that the same participant made during the Go trials (each circular panel represents the distribution of movement for each of the four targets during the plan repetition block).

The No-Go trials allowed us to dissociate planning from execution of a movement, and biasing a specific target allowed us to manipulate the probability distribution of upcoming plans toward that target, and then test the effect of this manipulation on physiological biases. Yet, one might be concerned that having many No-Go trials, in which participants did not make any actual movement, might discourage participants from truly planning any movement. If this was true, we would expect increase or unchanged reaction time in the later trials of the plan repetition block. Instead, if participants repeatedly planned the desired movement, we would expect a selective reduction in reaction time for the frequent target. Our data revealed a significant time effect (Early vs. Late) (*F*_(1,9)_ = 30.09, *p* = 0.004), and a significant interaction between the direction (repeated plan vs. other) and time (Early vs. Late) (*F*_(1,5)_ = 20.8, *p* = 0.0061 Fig. 3G). Post hoc analysis revealed that, late in the plan repetition block, the average RT across all participants in the direction of the repeated plan was significantly lower than early in the repetition block (*t*_(9)_ = 6.98, *p* < 0.0001), and was significantly lower (*t*_(9)_ = 4.66, *p* = 0.0055) than the average RT in the non-frequent target directions. The average RT in the other targets decreased (*t*_(9)_ = 2.19, *p* = 0.045). Our data rule out the possibility that participants did not truly intend to execute the correct plan during the No-Go trials. Altogether, our data clearly show that physiological biases evoked from the human motor cortex do not reflect a readout of ongoing planning-related processing of goal-directed voluntary movement.

## Discussion

In a set of neurophysiological and behavioral paradigms, we systematically uncoupled prior history of movement executions and planning for upcoming movement and showed that physiological biases evoked by TMS over the human motor cortex are due to effect of execution-dependent prior history and not to history of the most probable plan of action, suggesting that movement repetition induced changes in the state of the neural activity within the motor cortex that can then be read out by non-invasive transcranial stimulation. Our approach was to manipulate the statistics of movement history and the statistics of plans for upcoming movements independently, and then to test the effect of each manipulation on physiological biases. Although at baseline physiological biases shared similar low-level movement kinematics (i.e., direction) with plan-based biases, these processes diverged later when the statistics of either movement history or plan history were altered.

Use-dependent plasticity typically refers to the cortical reorganization of the effector (e.g., thumb) representation in the human motor cortex following practice^28–30^. Subsequent to a short period of training, consisting of simple, voluntary and repetitive thumb movements in a specific direction, non-invasive stimulation exhibits the reorganization of the cortical representation of the thumb by eliciting movement that encodes the low-level kinematic details of the newly practiced movement (i.e., bias toward the practiced direction)^28–30^. These findings suggest that repetition of simple movements leads to rapid establishment of a transient history-dependent change in the cortical motor network representing preferred thumb movements^1^. Long-term practice over years seems to have a similar use-dependent effect. TMS over the motor cortex in expert musicians (pianist & violinist) is more likely to elicit the same hand movements that occur while musicians actively play their instrument, compared to non-musician controls, or musicians that play other instruments^31^.

Multiple other factors might also affect physiological biases such as reinforcement^3^ and motor learning^31,32^. Observations from recent reports show that repetition of successful movements in the face of perturbations elicits stronger physiological biases than movement repetition alone^32–34^. Additionally, evidence from human neuropharmacology studies showed that administration of levodopa, which increases the presynaptic availability of endogenous dopamine, increased the strength of the biases evoked by TMS toward the repeated direction compared to a control group that only repeated the movement. This enhancement in physiological bias was correlated with the dopamine released in the striatum^27,35^. These results may suggest a hidden process, likely driven by success-related reinforcement signals that might modulate physiological biases. In addition, we have recently shown in a series of TMS experiments, that the physiological bias effect can be augmented if reinforcement is coupled with successful goal achievement during skill learning^36,37^.

However, what has not been clear, is whether these physiological biases might also reflect changes in planning-based processing that is related to an upcoming action. When hundreds of simple movements are repeatedly planned and executed in the recent past, it is possible that neural activity associated with the practiced movements becomes biased, not only toward the most frequently executed movement, but also toward the most frequent plan of upcoming movement^17^. Although not directly tested in TMS paradigms, this hypothesis was recently supported in goal-directed behavioral experiments by showing that a larger component of the behavioral bias primarily originates from changes associated with motor planning, with a second minor component that is mostly dependent on execution history^17,18^. Our findings revealed that physiological biases in motor cortex are insensitive to ongoing neural activity associated with the movement that participants are most likely planning to make next (i.e., the most frequent plan used in recent history), likely ruling out the functionality of evoked physiological biases with respect to voluntary goal-directed motor behavior^38^.

Although manipulation of plan history by changing the distribution of the locations of presented targets to include a more frequent target, thus biasing plans toward the more frequent target, significantly affected default plan for voluntary movement, it did not significantly affect physiological bias. Our result is in line with a recent neurophysiological study that recorded neural activity from M1 and the dorsal premotor cortex (PMd) while monkeys reached to randomly placed targets with different probability distributions of possible upcoming targets, and found that PMd activity, not M1, represents probability distributions of plan-based upcoming reaches, suggesting that such distributions are incorporated by the planning areas of the premotor cortex, outside M1, when coordinating goal-directed voluntary movement^39^.

Using a modified version of a delayed reaching task with No-Go trials allowed us to dissociate planning from execution of a movement. Repeating No-Go trials, thereafter, allowed us to bias the probability distribution of upcoming plans toward a specific plan. Nevertheless, having many No-Go trials, in which participants did not make any actual movement, might discourage participants from truly planning to execute the desired movement. This concern might influence the amount of planning and thus weaken the conclusion that TMS evoked-movement truly does not reflect the most probable upcoming plan. If this was true, we would expect increased, or unchanged, reaction time in the Go trials later in the planning repetition block. Instead, if participants repeatedly planned the desired movement, we would expect a selective reduction in reaction time for the frequent target. The significant time, as well as target × time interaction, effect supports the idea that participants repeatedly planned the desired action.

Our findings show that the dependency of physiological biases on execution history, and the insensitivity of these biases to prior planning, is analogous to some degree with the execution-dependent component of the behavioral biases found in goal-directed voluntary tasks^17^. Although the consistency between the two observations indicates that physiological and behavioral/movement biases might reflect similar underlying execution-dependent mechanisms, there are several factors that weaken this interpretation and even question the relevance of physiological biases to normal behavior. *First*, the strength of physiological biases is ten-fold greater than that of behavioral biases. On average, the magnitude of physiological biases is within the range of 150-180 degrees from baseline distribution of thumb direction^1,2^, whereas behavioral biases, at most, have a magnitude of 10 degrees^3^. *Second*, the timescale of physiological biases seems to be longer than behavioral biases. It was shown that TMS-evoked biases in thumb movement could last up to ∼40 min following 30 min of repetitions in the biasing direction. On the other hand, behavioral biases last at most for several trials (*n* = 20 *−* 50 trials) before drifting back to baseline level. *Third*, in contrast to physiological biases, behavioral biases appear to be task sensitive. During passive tasks, when a robot moves the hand to a target, behavioral biases were also observed^3^. However, when the thumb was passively and briskly moved in a direction opposite to the baseline TMS-evoked movement direction for 30 min, subsequent TMS did not reveal any physiological biases toward the repeated direction^40^. On the basis of these reports and recent observations^41^, we speculate that physiological and behavioral biases might coexist and share similar characteristics in some task setups, but they probably differ in the underlying mechanism. Future work is needed to directly examine the mechanism(s) that underlies each process.

At the neural level, our data can be reconciled with the recent theory of neural dynamic system. This approach posits that neural dynamics of population activity can be parameterized by a state equation that has different initial conditions and evolves over time. The evolved neural trajectory can span over multiple spaces, including separate, yet orthogonal, spaces for planning and movement execution^42^. The strength of the dynamical systems approach is that it provides a simplified mechanistic basis for understanding the link between time-varying activity of neural populations and planning and execution motor behavior^42–44^. Within this dynamic system, it is believed that movement preparation involves setting the state of the motor cortex to a particular, movement-specific state^45,46^, where preparation is thought to set the initial state of a dynamical system that generates patterns of activity required for movement^43,47,48^. Our results showing that modulation of execution history significantly affected physiological biases might therefore be a consequence of altering the default state in the movement space in M1 for the repeated direction, with little effect on ongoing activity of the preparatory space (Fig. 4). Stimulation of the motor cortex may thus reflect a readout of the cortical state within the movement space where movements located close to the default state may be more likely to be elicited by TMS. Altering frequency of potential targets shifted the distribution of plan-based biases toward the frequent target, with no effect on physiological biases. This can be explained by changes in the default preparatory state of cortical activity that converged into a new default state in the preparatory space. For instance, when no target ever appeared or in regular trials with low RTs (i.e., ≤50 msec), requiring participants to move is likely to trigger a readout of the preparatory state of the motor system at that moment.

**Figure 4.**
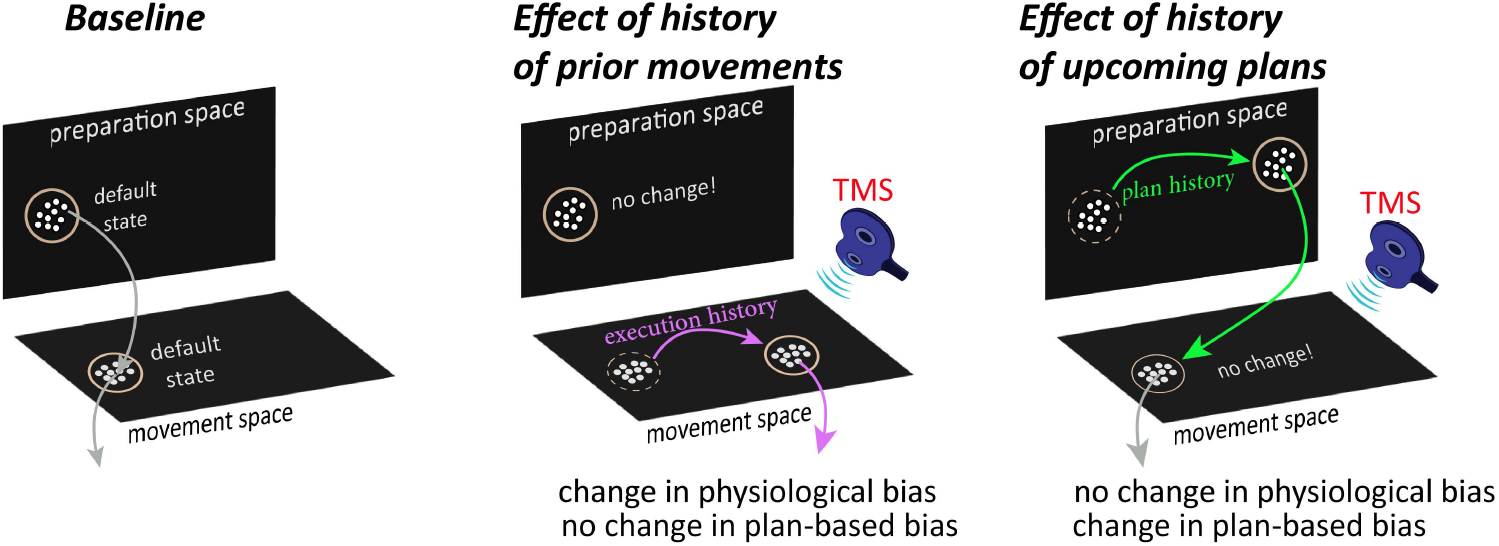
Proposed dynamic of neural activity in the motor cortex underlies physiological and plan-based biases. Recent execution history alters the default state in the movement-related space toward the repeated direction, with little effect on ongoing activity of the preparatory space. Stimulation of M1 may thus reflect a readout of the cortical activity where movements located close to the default state may be more likely to be elicited by TMS. On the other hand, manipulating plan-based expectation changed the default preparatory state.

The absence of representation of recent motor plans in the TMS-evoked movements does not necessarily mean that M1 cannot reflect functional changes associated with motor planning. In fact, a rich set of studies have now established the general finding that M1 corticospinal excitability is modulated during action preparation in an effector-specific manner^15,49–54^. Specifically, it has been shown that the corticospinal excitability (CSE), as measured by motor-evoked potentials (MEPs) of specific muscles following TMS over M1, allows for differentiating between various intrinsic muscles, and thus provides sufficient resolution to distinguish the physiological underpinnings of action preparation and selection for different finger movements^15^. However, there may not be a straightforward relationship between MEPs and motor output changes (as for example, evoked movement). MEPs can therefore indicate that something is changing physiologically during motor preparation, but the relationship of these measures to the changing behavior remains to be determined.

What exactly is probed in our catch trials when no visual target ever appeared? We believe that when requiring the participant to move on an imperative cue (i.e., the fourth audible tone) without the target being presented, the current state of the motor plan (i.e., default plan) becomes accessible and can be revealed via behavioral experimentation (e.g., catch trials). Before we discuss the potential cognitive-motor processes that might occur in catch trials, let us first consider a regular trial with sufficient RT when the target appeared at some point before the 4^th^ imperative tone. During the reaction time of these trials, a series of processes occurred that enabled the brain to perceive the surrounding environment, identify a particular target (or object) of interest, determine the required action in response to that target, and send a motor command to implement the desired action. These processes involve higher-level perceptual decision-making (e.g., processing sensory information to identify the location of the target), conception of the motor goal to be achieved, and movement-related processing (e.g., action selection for the thumb to achieve the goal) that may occur in parallel^11,55,56^. In catch trials, the motor goal is not well defined as the visual target never appeared, thus limiting the perceptual decision-making process for identifying target location. It is possible, however, that participants used strategic guess about which target would appear or strategic decision to move in the most comfortable or easiest direction. Nevertheless, our observation that the distribution of movement directions with very low RTs, in the timed-response trials when targets did appear, is very similar to that of movement in catch trials, suggests that the motor plan was successfully probed in these trials. Factors such as “comfort” and “ease” (e.g., minimization of energy expenditure, minimization of opposition from mechanical constraints such as tendons and ligaments, etc.) may constitute part of the “default plan” for choosing how an action will be generated to achieve a goal, as may be the case in low RT trials and catch trials. At this stage, we cannot resolve the “default plan” into its constituent components or completely disentangle the influence of higher-level cognitive planning from movement plan *per se*, and further work, beyond the scope of the present investigation, will be required to fully understand the relative contribution of each component of the cognitive-motor processes.

Taken together, the present results may imply that physiological change of the effector representation in the motor system incorporates an abstract representation of the kinematic structure of the executed behavior of recent prior movement. These physiological markers of change, however, were not indicative of ongoing activation that is related to planning of upcoming actions.

## EXPERIMENTAL PROCEDURES

### Participants

A total of 40 young participants (22 females; age 24.31±3.44 years; mean±s.t.d.) were recruited to participate in three experiments. All participants were healthy and right-handed. Participants provided written consent to participate in the study, which was approved by the institutional review board and ethics committee of the Technion-Israel Institute of Technology. Participants were given monetary compensation for their participation (100 ₪≈ $30).

### Apparatus

The experimental protocol involved goal-directed and TMS blocks in different sequences for the three experiments (see Procedures section below, and Fig.1C, 2A, 3A) in order to dissociably manipulate history of movements and plans, while probing physiological and/or plan-based biases before, during, and/or after. In the TMS neurophysiological sessions, participants were seated in a chair and were firmly connected to a frame that kept the head steady and the stimulating coil in a constant position with respect to the head. Each participant’s dominant forearm was immobilized using a customized armrest, with the four long fingers supported, and the thumb entirely unconstrained and free to move. Single TMS pulses that evoke involuntary thumb movements were delivered over the motor cortex. The point of TMS application is considered optimal if thumb movements are evoked in a consistent direction with stimulus intensities slightly above the movement threshold^1,2^. In goal-directed behavioral motor task sessions, participants were asked to make fast voluntary center-out ballistic movements, with the same thumb, toward 5-mm diameter targets presented at an equal distance of 4 cm from the starting point (Fig. 1A-B). Three TMS-behavioral experiments were undertaken to address the study aims.

#### TMS neurophysiological paradigm and apparatus

TMS sessions were performed using a PowerMag stimulator (The Mag & More®, Germany) with the participant at complete rest, defined as absence of visible or audible background electromyography (EMG using Delsys^®^, USA) activity exceeding the noise level of 25 μV and confirmed by measuring surface EMG activity from abductor pollicis brevis muscle (APB) and flexor pollicis brevis (FPB) muscles (agonist and antagonist muscles, respectively) of the right hand, both known to be activated during thumb movements^1,2,57^. A 70-mm figure-of-eight magnetic coil was placed tangential to the scalp with the handle pointing backward and laterally at a 45° angle away from the midline, approximately perpendicular to the central sulcus. To ensure accurate and consistent positioning of the TMS coil throughout experimental sessions, a frameless 3D neuro-navigation system (The Mag & More^®^, Germany) was used. The optimal scalp position (i.e., hotspot) for activation of the APB was identified using a stimulus intensity sufficient to evoke small thumb movements. In this optimal spot, the resting motor threshold, a measure of neuronal excitability, was determined as the minimum TMS intensity that evoked motor-evoked potentials (MEPs) of ≥ 50 μV in 5 out of 10 trials at rest^58,59^. The intensity of TMS for eliciting isolated and mild thumb movements in a consistent direction was set above the resting motor threshold. The direction and amplitude of TMS-evoked thumb movements were recorded using six motion-capture cameras (Prime 13, Optitrack^®^, USA) to record the trajectory of a spherical reflective marker placed on the thumb, at a sampling rate of 240 Hz. The position of the reflective marker was mapped onto a screen, in which the x and y axes represented thumb adduction-abduction and flexion-extension, respectively.

#### Goal-directed behavioral paradigm and apparatus

The participant’s dominant forearm was restrained in a molded armrest in a semi-pronated position, such that the four fingers were in a slightly extended position and the thumb was entirely unconstrained. The use of the armrest allowed consistent positioning of the arm and hand across the different sessions of the experiments. The same motion-capture camera setup as in the TMS paradigm was used to record thumb trajectory. The participant was able to control a screen cursor by moving her/his thumb (Fig. 1B). Participants performed three types of movement trials (each in a separate block): *timed-response, movement repetition*, and *plan repetition* trials. In *timed-response* trials (Experiments 1-3), after entering the central start position, participants heard a sequence of four tones, spaced 500 ms apart. One of four potential targets appeared 50-300 ms prior to the fourth tone, and participants were instructed to move to the target as quickly as possible synchronously with the onset of the fourth tone^20–22^. Manipulation of the timing of the target appearance allowed us to effectively impose a relative RT on each trial and thus control available response preparation time. We characterized the dynamics of response preparation for each target by assessing the accuracy of participants’ movements as a function of the imposed RT. The predictable sequence of auditory tones serves to minimize the ambiguity about the time of target presentation, which is known to increase RTs^20,23^. Movement initiation time was determined online as the time at which the tangential velocity first exceeded 5% of maximum velocity. If participants failed to initiate their movement within ±150 ms of this time, an on-screen message was shown, “Too Early” or “Too Late”, accordingly. The timed-response trials included a subset of catch trials in which no target ever appeared, and participants were required to move with the onset of the fourth tone to any direction of their choice. These catch trials enabled us to assess participants’ default preparation in the absence of a specific presented target location (see Discussion about the interpretation of catch trials). With respect to these trials, participants were instructed to move as quickly as possible to any direction when they heard the 4^th^ tone if no target appeared. In the timed-response trials in general, participants were not given any specific instructions or information about potential targets (e.g., time of target presentation, locations, etc.). Inter-trial interval (ITI) in the timed-response task was set at 2 seconds. In *movement repetition* trials (Experiment 2), participants were instructed to repeat the same movement toward the center of a semi-circular arc target. The center of the arc was 180 degrees from the baseline TMS-evoked direction. Participants were instructed to make quick and accurate movements following presentation of the arc. The *plan repetition* trials were similar to the *timed-response* trials, but implemented a Go/No-Go cue as well as an altered target probability distribution (i.e., 70, 10, 10, 10%) and consisted of 750 trials. Additionally, a single TMS pulse was delivered during some of these trials, 150 msec before the Go/No-Go cue (see Procedure section, Experiment 3 for details).

### Procedure

#### Experiment 1

The experiment was designed to determine whether baseline TMS-evoked physiological biases share similar low-level movement kinematics with plan-based biases of voluntary movement estimated using the timed-response task. Participants (n=15, aged 24.3±2.3 years) performed three consecutive blocks (TMS-behavioral-TMS; Fig. 1C). In the first block, participants underwent a TMS paradigm. During this block, direction of the thumb movement was established by delivering a series of 65 TMS pulses at 0.2 HZ to the optimal scalp position that evoked isolated involuntary thumb movements. The second block included a goal-directed task. Participants were asked to perform 150 timed-response thumb movements, either toward a target (120 trials), selected randomly from four targets (*θ*_*base*_+ [0°,90°,180°, and 270°]) positioned 4 cm from the origin, or toward the direction of their choice in the absence of a presented target (30 trials). Targets were presented randomly 30 times each. Finally, participants again underwent a TMS block identical to the first TMS block. The 2^nd^ TMS session was conducted to verify that short practice of timed-response goal-directed tasks is insufficient to elicit physiological biases^1,27^, such that baseline measurements would be distinct and not confounded by other biases. Time between blocks was ∼3-5 seconds.

#### Experiment 2

In this experiment, we systematically manipulated the probability of recent history of executed movements by asking participants to repeatedly make thumb movements toward a novel direction located in the opposite direction of each participant’s baseline TMS-bias (*θ*_*base*_). We estimated the physiological biases using TMS, and plan-based movement biases using the catch trials of the timed-response block, before and after participants voluntarily repeated a movement toward the center of a semi-circular arc target set at 180° from *θ*_*base*_. Specifically, participants (n=15, aged 25.2±4.3 years) were asked to perform three consecutive sessions: *baseline, movement repetition* and *post-movement repetition* (Fig. 2A). The *baseline* session included one TMS block and one goal-directed behavioral block, as in Experiment 1. In the *movement repetition* session, participants were instructed to repeat the same thumb movement for 40 min toward the repeated direction, opposite to that evoked by the *baseline* TMS session (*φ*_*rep*_ = *θ*_*base*_ *−* 180°). In the *post-movement repetition* session, participants underwent another TMS block and another goal-directed block.

#### Experiment 3

To further test the relationship, or lack thereof, between physiological biases and ongoing activity related to motor planning, we systematically dissociated prior history of movement execution from prior history of movement plans. To do so, we biased the probability of upcoming movement plans by making potential target locations (identical to those in the timed-response task) not equally probable and, instead, increased the likelihood of one of the four target locations, while implementing a Go/No-Go goal-directed paradigm. Participants (n=10, aged 22.7±2.9 years) were asked to perform three consecutive sessions: *baseline, plan repetition*, and *post-plan repetition* (Fig. 3A). The *baseline* session included one TMS block and one goal-directed behavioral block, as in Experiments 1&2. The *plan repetition* session included a goal-directed behavioral block with unequal target location probabilities. For each participant, the frequent target was set in the direction with lowest probability of the fitted von Mises distribution for the movement direction in catch trials during the baseline timed-response block. Unlike the baseline block in which the four targets were uniformly distributed, the frequent target was presented in 70% of trials (525 trials), whereas the other three were each presented in 10% of trials (75 trials per target). A Go/No-Go paradigm was implemented such that in 80% of trials participants were led to prepare a movement, but were cued thereafter not to execute it (No-Go cue), and in 20% of trials participants were cued to execute the planned movement (Go cue). The target appeared 750 msec before the Go/No-Go cue, which was indicated by the gray target either turning green or red, respectively, simultaneously with the fourth tone. Within the *plan repetition* block, TMS was implemented, starting after the participant completed 500 trials of *plan repetition*, to probe change, or lack thereof, in physiological bias due to plan history manipulation, and was delivered at random in 15 trials for each target, 150 msec before the Go/No-Go cue. A *post-plan repetition* block was performed following the *plan repetition* block, which consisted of a modified timed-response block, integrating TMS to test physiological biases within the same context as probing of any changes in the default plan resulting from *plan repetition*. Thus, the *post-plan repetition* block consisted of an identical timed-response task as the previous ones, but with a single pulse of TMS delivered in 15 out of the 30 catch trials, 150 msec before the fourth tone.

### Data Analysis

Physiological TMS and behavioral data were analyzed using custom routines in Matlab to compute physiological biases, plan-based biases, repeated direction, reaction time, accuracy, and speed-accuracy trade-off function.

#### Physiological biases (i.e., TMS-evoked)

The direction and amplitude of TMS-evoked thumb movements were determined using six motion-capture cameras that recorded the trajectory of a spherical reflective marker placed on the thumb, at a sampling rate of 240 Hz. The position of the reflective marker was mapped onto a screen, in which the x and y axes represented thumb adduction-abduction and flexion-extension, respectively. Given the x and y position of the marker, the velocity of the involuntary thumb movement was calculated. The thumb direction was defined as the direction at movement onset. Movement onset was specified, offline, as the timepoint when the velocity of the cursor exceeded 5 % of peak velocity. Then, distribution of thumb direction was estimated using the von Mises distribution^24^, and the direction with highest probability of the fitted von Mises distribution was considered as the physiological bias.

#### Plan-based biases (i.e., default plan)

During catch trials, no target ever appeared and participants were required to move with the onset of the fourth tone to any direction. These catch trials enabled us to assess participants’ default plan in the absence of a specific presented target location. The distribution of thumb direction in these catch trials was estimated using the von Mises distribution, and the direction with highest probability of the fitted von Mises distribution in catch trials was considered as the plan-based bias. After defining the plan-based direction, we defined the target near the plan-based bias (near-PB) in the regular timed-response trials as the closet target with minimal distance (≤ 45°) from the direction of the plan-based bias. All other targets were defined as the far-PB targets.

#### Reaction time (RT)

During the timed-response trials, RT (or effective RT) was calculated as the time interval between target presentation and movement onset. During the Go trials of the plan repetition block, RT was calculated as the time interval between Go cue and movement onset.

#### Speed-accuracy trade-off function

For each target, the probability of initiating an accurate movement (the *success rate*) in the timed-response trials at any given RT was estimated based on the proportion of accurately initiated movements within a 125-ms window around that RT. A movement was considered accurate (i.e., successful) if the initial direction of the movement was within ±45° of the target direction. Otherwise, the movement was considered an error. This yielded an estimate of the speed–accuracy trade-off function (Supplementary Fig. S1B). In order to quantitatively characterize this trade-off, we followed the likelihood model presented by previous studies^20,60^, in which a single preparation event is assumed to occur at a stochastic time 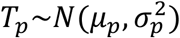, with movements initiated before *T*_*p*_ directed randomly, and movements initiated after *T*_*p*_ directed accurately toward the target. These assumptions lead to the speed-accuracy trade-off following a cumulative Gaussian shape (Supplementary Fig. S1C). Since we fit this model to the different targets, participants may have expressed preferred default movements at very low RTs and would thus have been either above or below chance. We allowed for this possibility through a parameter *α*_0_ that defined the participant’s lower limit accuracy. We also allowed for the fact that participants were not 100% accurate even at long RTs through a parameter *α*_∞_ that defined the participant’s asymptotic accuracy.

We estimated these parameters (µ_*p*_, 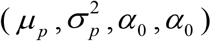, *α*_0_,*α*_0_) based on maximum likelihood. The probability of the movement in trial *i* being accurate (*H*^*i*^ = 1) given that it was initiated at time *RT*^*i*^ was then given by:

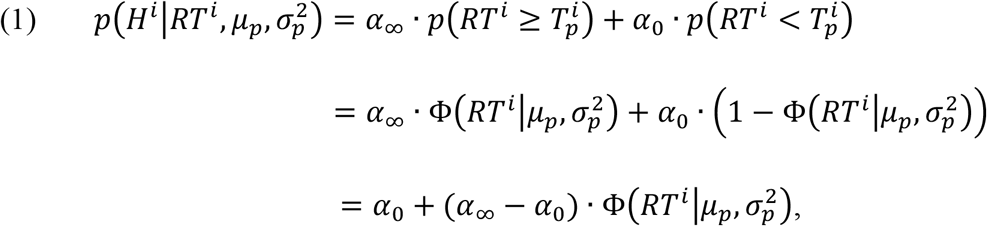

where 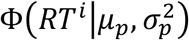 is the cumulative normal distribution function. The log-likelihood function for each trial was therefore defined as:

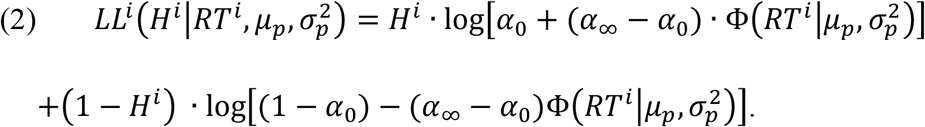

The total log likelihood of the parameters *α*_0_, *α*_∞_, *µ*_*p*_, *σ*_*p*_ was therefore given by:

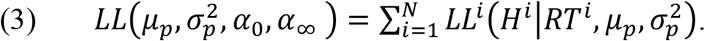

In order to prevent overfitting and to improve the generalization of our model, we used a regularization technique, introducing a penalty on the variance (i.e. 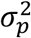) of *T* as follows:

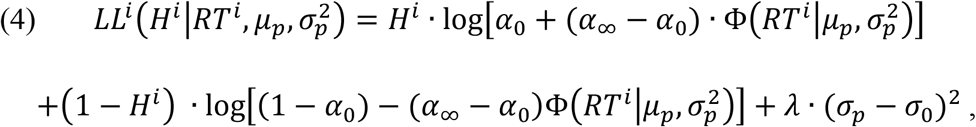

where *λ* is the regularization term, which we fixed at *λ* = 1000, and *σ*_0_ corresponds to a slope prior, which we set at *σ*_0_ = 0.06. Parameter estimates were obtained separately for targets near the plan-based bias direction and for targets far from the plan-based bias direction based on the pooled data from all participants in the timed-response condition. We used the pooled data because the data for individual participants was too sparse to obtain reliable fits. Since we were interested in comparing the accuracy at low RTs near the plan-based bias direction vs. the other far plan-based bias targets, we collapsed together the data of all directions toward targets far from the plan-based bias direction. We calculated confidence intervals for parameter estimates through a bootstrap analysis, using 1,000 unique combinations drawn with replacement from the subject pool. The 95% confidence intervals were calculated as the 2.5 and 97.5 percentile values from the distribution for each coefficient obtained across the 1,000 fits. To visualize the distribution of reach directions in the catch trials in Experiments 1, 2 and 3, we used a kernel density estimation procedure using von Mises kernels with a kernel width of 30°.

### Statistical Analysis

The statistical analysis was performed using the Circular Statistics Toolbox (Directional Statistics) of Matlab software (MathWorks) and Prism software (GraphPad). A two-sample Kuiper’s test^61^ was used to determine the similarity, or dissimilarity, between the distribution of physiological biases and plan-based biases at baseline. Kuiper’s test is invariant under cyclic transformations of the independent variable. This invariance under cyclic transformations makes Kuiper’s test valuable when testing for differences between circular probability distributions. In the Circular Statistics Toolbox, this test can be performed using the *circ_kuipertest*() function. To test the effect of manipulating statistics of history of prior movement (Experiment 2) or history of upcoming plans (Experiment 3), we ran Kuiper’s test on the change (i.e., *post-pre* training) of physiological biases and plan-based biases and compared the distribution of the deltas to the normal distribution with *µ* = 0, *σ* = 10°. We also used Kuiper’s test to determine whether the direction of movements occurring before, and immediately after, the target was displayed were comparable to the plan-based directions in the catch trials during the timed-response task. Of note is that the *p value* of Kuiper’s test is taken from tabulated values determined previously^61^. In this table, any value of *p* > 0.1 is set to *p* = 1. To determine the accurate value in all cases when *p* > 0.1 (not significant), we used the equivalent one-sample *t − test* for circular data and tested whether the difference of angles was significantly different from normal distribution with *µ* = 0, *σ* = 10°. Lastly, to assess the accuracy at low RTs of the speed-accuracy trade-off in Experiments 1, 2 and 3, we ran a bootstrap analysis on the parameters *α*_0_ that defined the participant’s lower limit accuracy, with 1,000 resamples (for details, see Data Analysis above). In all comparisons, the significance level was set at 0.05.

## Acknowledgment

We would like to thank Adrian Haith and Mona Khoury-Mireb for the helpful comments and discussions. This study was funded by the Israel Science Foundation (ISF) Grant 1634/19 (FM) and the United States - Israel Binational Science Foundation (BSF) Grant 2021323 (FM).

## Supplemental Information

### Cortically-evoked movement in humans reflects history of prior executions, not plan for upcoming movement

Abdelbaset Suleiman, Deborah Solomonow-Avnon and Firas Mawase

**Figure S1.**
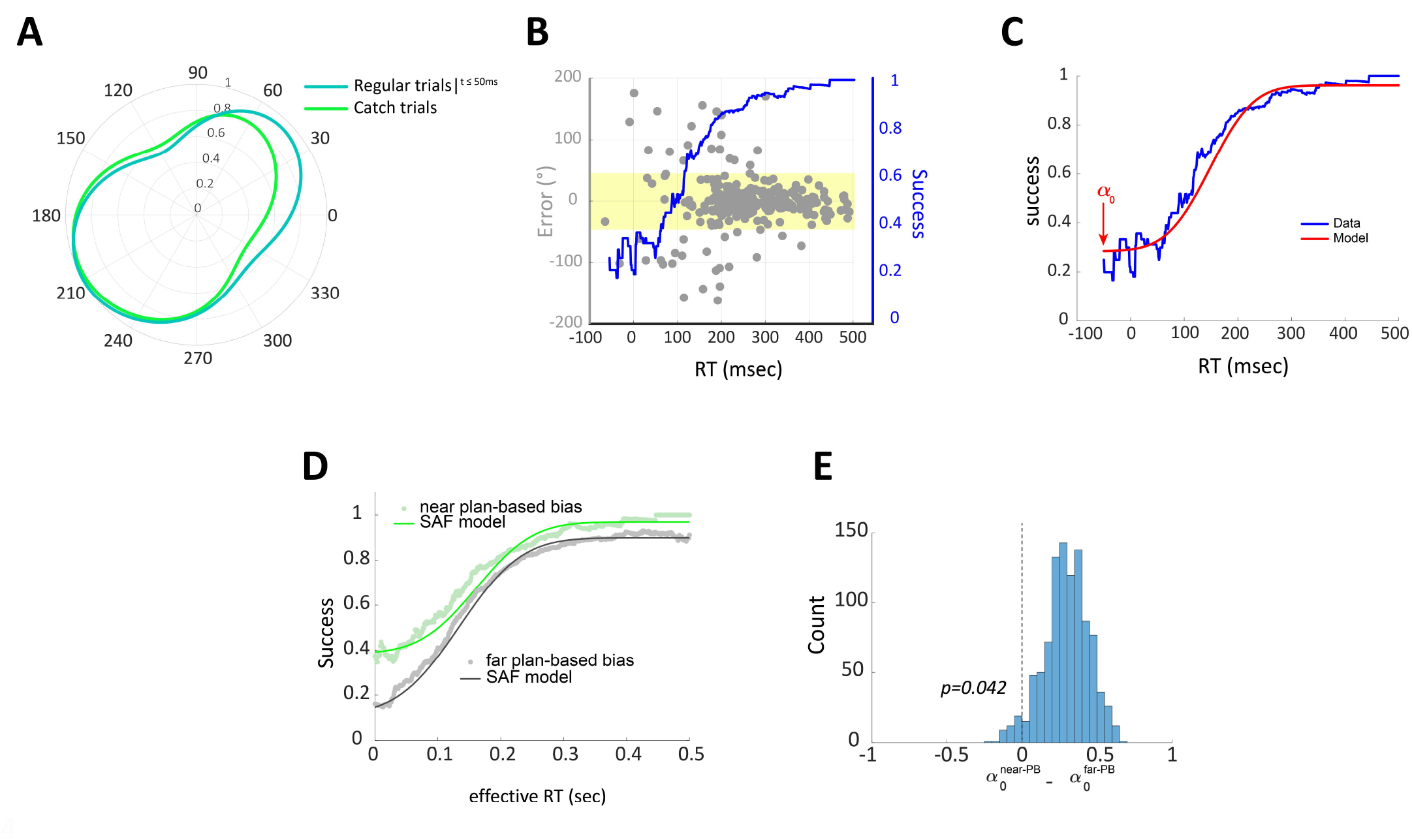
Estimating Speed-accuracy trade-off function and confirmatory analysis of default plan in catch trials and trials with low RTs in Experiment 1. **A**. The distribution of movement directions in the timed-response trials occurring any time before the target appeared, and up to 50 msec immediately after the target was displayed (dark green) were comparable to the directions in the catch trails (light green). **B**. Example of data during the timed-response condition pooled across participants in Experiment 1 (target #2). Gray points indicate direction error (°) as a function of RT for individual trials; blue line shows the moving average of the probability that a movement is successful for a given RT, which corresponds to the speed-accuracy trade-off. Yellow area indicates the range of where a given movement was considered successful. **C**. Illustration of maximum likelihood model fit (red line) to empirical speed-accuracy trade-off data (blue line). Arrow indicates the estimated parameters *α*_0_ that defined the participant’s lower limit accuracy. **D**. Speed-accuracy trade-off for movements near the plan-based direction revealed that this movement involved the default plan. This was reflected by the increased accuracy at short reaction times (RTs) for targets near the plan-based bias, but not for the other targets. **E**. Fitting the speed-accuracy function revealed significant difference of the parameter *α*_0_ (which reflects participant’s lower limit accuracy) between near plan-based (near-PB) target and the other targets (far-PB).

**Figure S2.**
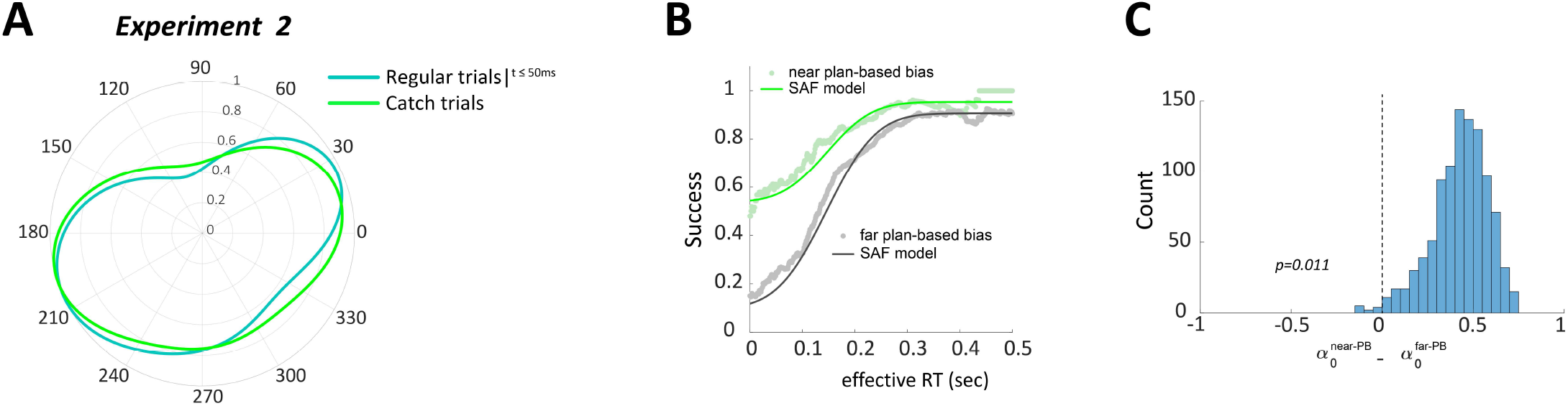
Confirmatory analysis of default plan in catch trials and trials with low RTs in Experiment 2. **A**. The distribution of movement directions in the timed-response trials any time before the target appeared, and up to 50 msec immediately after the target was displayed (dark green) were comparable to the plan-based directions in the catch trials (light green). **B**. Speed-accuracy trade-off for movements near the plan-based direction revealed that this movement involved the default plan. This was reflected by the increased accuracy at short reaction times (RTs) for targets near the plan-based biases, but not for the other targets. **C**. Fitting the speed-accuracy function revealed significant difference of the parameter *α*_0_ (which reflects participant’s lower limit accuracy) between near plan-based (near-PB) target and the other targets (far-PB).

## Notes

**Conflict of Interest:** The authors declare no competing financial interests.

### Competing Interest Statement

The authors have declared no competing interest.

